# Within-species control reveals novel immune response genes in *Caenorhabditis elegans*

**DOI:** 10.1101/2022.05.31.494203

**Authors:** Ayush Ranawade, Emma Hartman, Erel Levine

## Abstract

The nematode *Caenorhabditis elegans* is a simple model host for studying the interaction between bacterial pathogens and the metazoan innate immune system. In the last two decades, much focus has been given to studying intestinal infection in the worm by the clinical strain *Pseudomonas aeruginosa* PA14. Powerful genetic and molecular tools in both species facilitate the identification and analysis of bacterial virulence factors as well as host defense factors.

However, findings from these studies are confounded by the use of the genetically, metabolically, and physiologically divergent *E. coli* OP50 as a food source and as the non-virulent control. Here we report the use of *P. aeruginosa* z11 strain as a preferable control for PA14 infection studies. We demonstrate that many aspects of worm behavior, health-span, longevity, food attraction, brood size, lifespan, and pathogen avoidance and survival are not affected in z11. We show that the use of z11 as a control for transcriptomics analysis can increase the discovery power. In particular, we identified three novel pathogenesis-related genes in *C. elegans*. Our findings accentuate the importance of choosing an appropriate control environment for host-pathogen studies.

## Introduction

*C. elegans* has been a well-established model to study aging, development, genetics, transgenerational inheritance, and pathogenesis of diverse types ^1–4^. Several characteristics of *C. elegans* make it a valuable tool for studying these aspects of biology; for example, worms have a short life cycle, large fecundity, genetic homogeneity due to hermaphroditic reproduction, and a wealth of available genetics and imaging tools ^3,5^. Additionally, 60-80% of proteins of *C. elegans* have a homolog in humans, and 40-75% of disease-related genes are conversed, leading to surprisingly high potential for translational studies ^6^. Remarkably, mutations in these homologous genes in *C. elegans* produce quantifiable phenotypes that are utilized to understand the molecular mechanism of the disease-associated genes and high throughput drugs screenings ^7^.

In the last two decades, *C. elegans* has been used as a host model for studying infection by diverse types of pathogens, including bacteria, fungi, viruses, and more ^8^. One of the best studied pathogens is *P. aeruginosa* PA14 (PA14 hereafter), an opportunistic pathogen isolated from cystic fibrosis patients and known to cause sepsis in burned patients, pulmonary infections, and mortality in cystic fibrosis patients ^9^. This environmentally persistent pathogen can employ diverse virulence factors that allow PA14 to be highly infectious to a broad range of hosts, including *C. elegans* ^10^. The *Pseudomonas* studies in *C. elegans* have led to identifying the mechanism of innate immune response pathways, pathogen avoidance behavior, and small noncoding RNA-based transgenerational memory ^11,12^. For these studies, *Escherichia coli* strain OP50 (OP50 hereafter) has been used as a reference strain: worms were fed OP50 during cultivation, and plates with *C. elegans* fed by OP50 were used as control against which all the experimental assessments were compared. However, OP50 and PA14 belong to different bacterial families, *Enterobacteriaceae* and *Pseudomonadaceae,* respectively, and exhibit significant genetic, physiological, and structural differences. For example, B-type *E. coli* strain OP50 is a facultative anaerobe, coliform, biofilm deficient, non-motile, non-pathogenic, uracil auxotroph that forms thin bacterial lawns ^13^. In contrast, PA14 is an obligate aerobe, non-coliform, biofilm-forming, motile, opportunistic pathogen. Furthermore, *C. elegans* have been shown to exhibit bacteria dependent lifehistory traits, including variations in development, lifespan, reproductive lifespan, metabolism, and pathogenicity ^14–19^. These phenotypic variations in *C. elegans* are associated with genetic, metabolic, and surface molecules cues from different bacterial strains ^20^.

In a typical study of the host response to a PA14, one measures some phenotype – survival, behavior, gene expression, neuronal activity, and more – at some time post-infection and compares it to the same phenotype observed in uninfected animals. The rationale of such an experiment is that observed differences are likely to be due to the pathogenic effect of PA14. However, given the many differences between OP50 and PA14 unrelated to pathogenesis, one must entertain other probable causes for the observed phenotypic differences. Moreover, some responses to pathogenesis may be masked by contradicting adaptation to other differences between these species.

In this study, we overcome these pitfalls by using *P. aeruginosa* strain z11 (z11 hereafter), a putative non-pathogenic strain, as a reference strain for *C. elegans* PA14 infection studies. The z11 strain was isolated from cystic fibrosis patients ^21^. A recent study that compared the lifespan of worms on different strains of *P. aeruginosa* found that worms have a similar lifespan on z11 as they do on a control *E. coli* strain HB101, suggesting the non-pathogenic nature of z11 ^22^. Here we show that worms raised on z11 demonstrate similar physiological and behavioral traits as those raised on OP50, with minor differences in developmental rate. We find that the transcriptome of worms exposed to the pathogenic PA14 is more similar to z11 than to OP50 but can also uncover important variations masked by the more divergent control, including identifying novel genes related to host response. These data sets represent a critical reference for evaluating studies of PA14 pathogenesis and underscore the importance of the microbial environment on the study and interpretation of host-microbe interactions.

## Results

### Z11 diet does not affect physiology, lifespan, and fecundity of C. elegans

It is well known that bacterial genotype and metabolism differentially affect worm physiology ^18,19^. We hypothesized that a non-pathogenic *P. aeruginosa* strain would be a better control for a *P. aeruginosa* PA14 infection than genetically divergent *E. coli* strains. We focused on the stain z11, which was reported to have no adverse impact on worm lifespan ^22^, while the morphology of its lawn, its antibiotic resistance, and its pyocyanin production suggest its proximity to PA14 (Supp. Fig. 1A). To establish z11 as a suitable non-pathogenic control, we first sought to determine that its effects on *C. elegans* physiology, development and behavior are similar to those on the well-studied benign strain OP50.

First, we asked if worms could be cultured on z11 as the sole source of nutrition. We maintained worms on z11 bacterial diet for >10 generations prior to performing a comparative physiological analysis. We then measured the lifespan of wild type N2 worms grown on these two species of bacteria. The worms grown on z11 at 25°C showed a slight extension in the mean lifespan of 15 ± 0.55 days compared to 14 ± 0.53 days for worms grown on OP50 (Fig. 1A).

**Figure 1.**
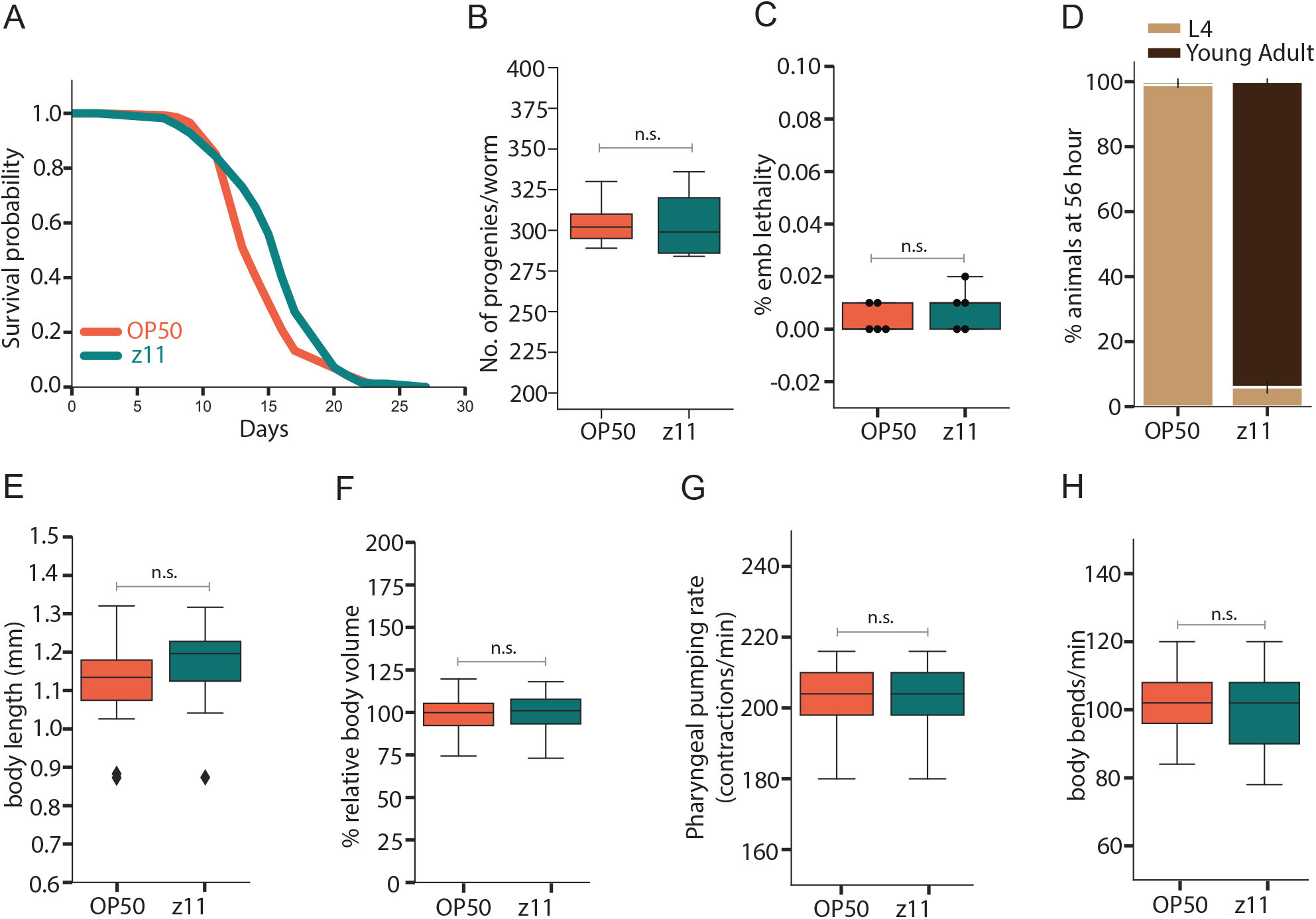
*P. aeruginosa* z11 does not affect life history traits of *C. elegans* (**A**) Survival of adult worms grown on OP50 (orange) and on z11 (teal) is not significantly different (log rank p-value 0.366). (**B**) Reproductive fecundity of worms grown on OP50 and z11 is very similar (n=10-14, p-value=0.493). (**C**) Embryonic lethality in worms grown on OP50 and z11 are very similar (n=150, p-value=0.198). (**D**) Worms grown on z11 develop faster than OP50. Percentage of late-stage larvae (L4, light brown) vs adult worms (dark brown) at 56 hours post L1 larvae arrest (n=90, P-value < 0.001). (**E-H**) Several physiological characteristics of worm are not affected by diet on OP50 and z11: (**E**) worm length (n=30, p-value>0.236); (**F**) Body volume (n=30, p-value>0.075); (**G**) Pharyngeal pumping rate (n=30, p-value>0.371); (**H**) Motility, estimated by worm thrashes (n=42, p-value>0.345). All graphs represent the combined results of 2-3 independent repeats. Hereafter, whiskers extend to the minimum and maximum values within 1.1-1.5 times the interquartile range. n.s.: non-significant.

Next, we looked for possible effects that cultivation on z11 diet might have on *C. elegans* development. We first measured the brood size and embryonic lethality and found no significant difference in the brood size between worms grown on z11 and worms grown on OP50 (Fig. 1B). Similarly, almost all embryos laid by worms grown on both species of bacteria were viable (Fig. 1C). To determine the developmental timings, we evaluated the time for age synchronized L1 larvae to reach the L4 larval stage. We found that L1 worms grown on the z11 bacteria showed an accelerated development and reached the L4 stage within 32 hours, 12 hours faster than worms fed on OP50. Notably, when the latter worms reached the L4 stage, most worms grown on z11 were already young adults (Fig. 1D). However, this accelerated developmental in z11 fed worms did not affect other physiological characteristics of the adult worms. For example, we found that the worms grown on OP50 caught up in size by young adulthood and showed similar body length and size (Fig. 1E-F).

To further characterize the food-dependent physiological characteristics, we measured the pharyngeal pumping rate (Fig. 1G) and thrashing movement (Fig. 1H) of adult worms. Both showed no significant difference between worms grown on z11 and worms grown on OP50.

### Z11 does not induce immune response and avoidance behavior in C. elegans

Our next goal in establishing z11 as a non-pathogenic control was to ask if worm response to z11 mimics known responses to pathogenic bacteria. Exposure to pathogenic bacteria elicits specific gene expression changes. In particular, exposure to PA14 is known to robustly induce the expression of the immune response gene, *irg-1* (infection response gene 1) ^23^. This strong induction is specific to virulent PA14 and is not observed with other pathogens. Moreover, studies of an *irg-1::GFP* reporter gene showed that the strength of induction is directly correlated with virulence, where the most virulent strains induce the strongest expression ^24^. We measured the activation of the *irg-1::GFP* transcriptional reporter in worms grown on z11, OP50, and PA14. While *irg-1::GFP* transgene was robustly induced by infection with PA14, transgenic worms grown on OP50 and z11 did not display a detectable GFP expression (Fig. 2A,B). This supports the idea that z11 does not induce an immune response typical to the response to PA14.

**Figure 2.**
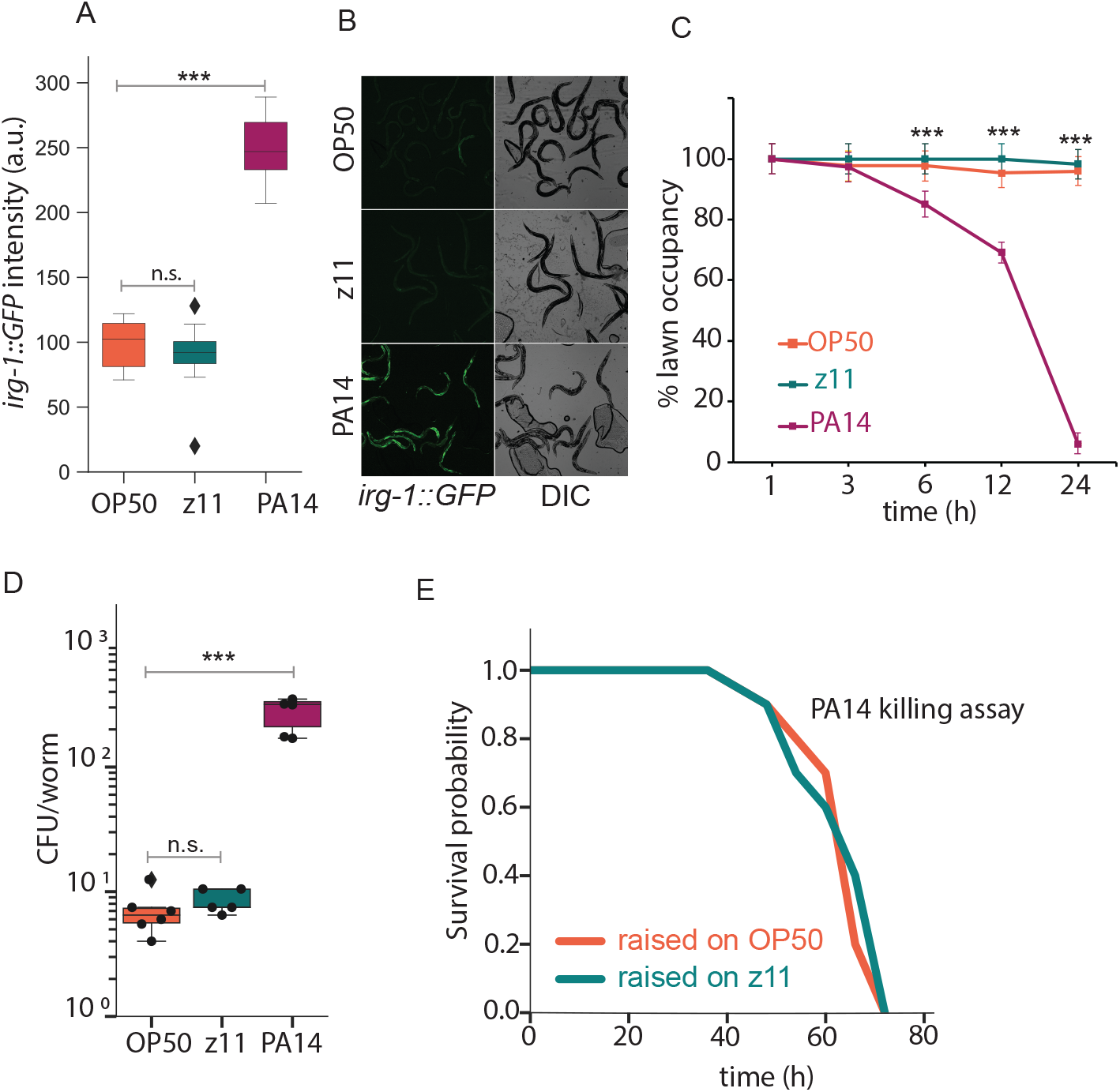
*P. aeruginosa* z11 does not induce immune response and pathogen avoidance in *C. elegans* (**A, B**) The fluorescent immune reporter *(irg-1::GFP)* is strongly induced by PA14 but not by OP50 or z11 (**A**) Quantification of GFP expression in arbitrary unit (a.u.). Hereafter *** p<0.001, n.s., non-significant. (**B**) Representative fluorescence and corresponding DIC images of each condition. (**C**) Worms learn to avoid the lawn of P14, but not the lawn of z11 or OP50. (**D**) Bacterial colony forming unit (CFU) obtained from the guts of animals grown on OP50, z11, and PA14 lawns. Worms show significant accumulation of gut bacteria on PA14 (n=60 p-value= 5.52422E-06) but not on z11 and OP50 (n=60, p-value= 0.021) E) Survival rates during a PA14 slow killing assay are the same for worms raised to young adulthood on OP50 or on z11 (log rank p-value = 0.456).

Worms are avid bacterivores who spend most of their time on the bacterial lawn when grown on the non-pathogenic OP50 ^25^. In contrast, when placed on a plate seeded with PA14 bacteria, worms are initially attracted to the bacterial lawn, but learn within 8 hours to avoid the pathogen by foraging off of the bacterial lawn ^26,27^. This pathogen avoidance behavior in *C. elegans* is directly related to the virulence of the bacterial strain, such that the less virulent PA14 *gacA* mutant fails to induce avoidance of the bacterial lawn ^28^. To test if z11 elicits pathogen avoidance, we performed an avoidance assay, where 30-40 young adults that were cultivated on OP50 are transferred onto a fresh bacterial lawn, and the numbers of worms on and off the lawn are counted over a period of 24 hours (Figure 2C). While worms remained on the z11 and OP50 lawns for the duration of the experiment, worms stated avoiding the PA14 lawn within 6 hours and were mostly away from the lawn by 24 hours. Similar results were obtained for worms cultivated on z11 for multiple generations (Figure S1B). This result demonstrates that z11 does not induce an avoidance response in worms that is typical to pathogen exposure and several abiotic stressors ^26,27,29^.

Worms can efficiently grind bacteria in the pharynx and lyse and digest them in the intestine ^30^. Pathogenic bacteria overcome these barriers and are able to accumulate in the gut ^31–33^. Consequently, young adult worms carry almost no live bacteria in their gut when exposed only to OP50 but accumulate a significant population of live bacteria in the gut when grown on PA14 ^32^. This prompted us to ask what is the size of the population of live bacteria in worms grown on z11. To address this question, we collected animals grown on OP50, z11, and PA14 and counted the number of colony-forming units (CFUs) per animal. We find that the gut population of z11 is similar in size to that of OP50, and significantly smaller than that of PA14, as expected (Fig. 2D). Given that (as described above) the rate of pumping z11 bacteria from the environment is similar to that of OP50, this result indicates that worms grind and lyse bacteria of these two species with similar efficiency.

Together, these results demonstrate that z11 is not pathogenic to *C. elegans* and does not induce genetic immune response or avoidance behavior.

### Cultivation on z11 does not alter the outcome of PA14 pathogenesis in wild type worms

The cultivation of worms on some bacterial species can protect them from subsequent infection by some pathogenic bacteria ^34^. For example, cultivation of worms on some isolates of *Lactobacillus* protects them from *Salmonella typhimurium* DT104 infection ^35^ while cultivation on the natural microbiota isolates *Pseudomonas lurida* MYb11 and *P. fluorescens* MYb115 protect the worm against pathogens such as *Bacillus thuringiensis* ^36^. In liquid media, worms grown on *E. coli* HT115 are resistant to infection PA14 and survive longer than worms reared on *E. coli* OP50 ^37^.

Because the metabolism of z11 is more similar to PA14 than OP50, we hypothesized that growth on z11 could better prepare worms for the transition from the cultivation food to PA14 and could therefore increase their survival. To test this hypothesis, we performed a slow killing assay as described previously ^10^ using worms grown on OP50 and z11. Surprisingly, we did not find any significant difference in the PA14 survival of wild-type worms grown on OP50 (54 ± 0.5 hours) and those grown on z11 (54 ± 0.55 hours) (Fig. 2E). These observations demonstrate that worm growth on z11 does not affect worm survival on PA14.

### The main difference between worm response to z11 and OP50 is metabolism; between response to z11 and PA14 it is proteolysis and extracellular transport

The data presented above demonstrate that z11 has no adverse effect on worm physiology or behavior, does not induce genetic or behavioral responses typical to unfavorable environment, and does not affect the dynamics of PA14 infection of the worm. Together these findings suggest that z11 can serve as a non-pathogenic control for worm infection assays.

To get a better understanding of what are common and distinctive effects of z11, OP50, and PA14 on the worm transcriptome, we performed RNA-seq of wild-type (N2) *C. elegans* fed on these three species for 12 hours and performed differential expression analysis between each pair of conditions. Below we call a gene differentially expressed if the difference between its expression in the two conditions exceeded 2-fold and with statistical significance p-adj values <0.01 (Table S1).

We first compared the expression of worms on PA14 or z11 with worms on OP50 (Fig. 3A). We hypothesized that some genes respond to some of the many differences between *E. coli* and *P. aeruginosa* and are therefore differentially expressed in both z11 and PA14, while other genes respond to the virulence of PA14 and are differentially expressed in PA14 alone. Indeed, 79% of the differentially expressed genes (DEGs) in the presence of z11 are also differentially expressed in response to PA14, while 71% of the genes differentially expressed in PA14 are *not* differentially expressed in z11.

**Figure 3.**
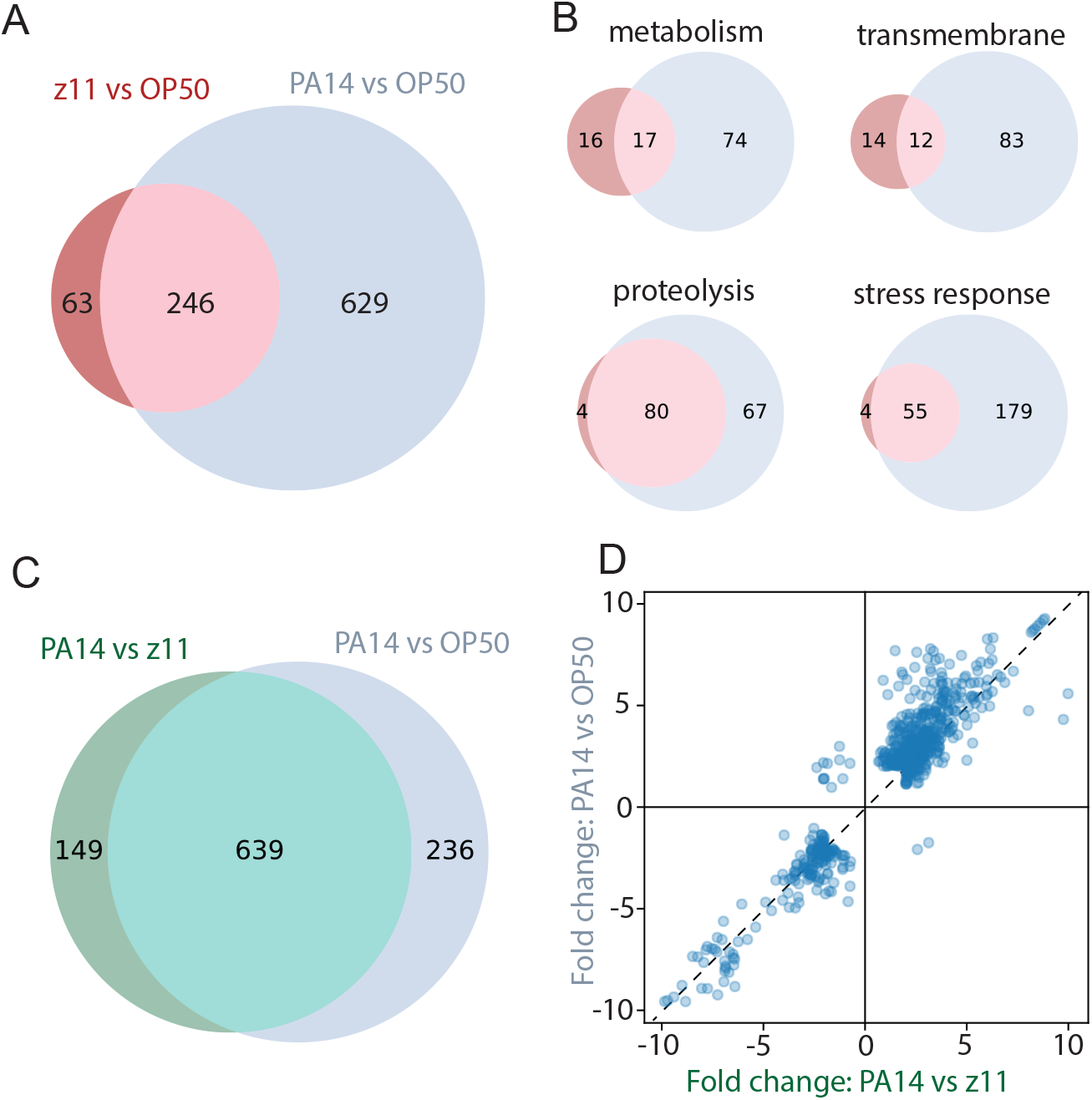
Bacteria-induced differential gene expression in worm reflects responses to diet changes and virulence (**A**) A smaller set of genes are differentially expressed between OP50 and z11 than between OP50 and PA14. (**B**) Genes associated with metabolism and transmembrane transport are more likely to be differentially expressed exclusively in either z11 or PA14, while genes associated with proteolysis and stress response are more likely to be differentially expressed in both. (**C**) Significant overlap between the genes differentially expressed between PA14 and z11 control and between PA14 and OP50 control. (**D**) High correlation (R^2^=0.92) in the expression fold-change of genes that are differentially expressed in both comparisons.

We used WormCat ^38^, a worm-specific tool for annotating gene-set enrichment data, to look for biological functions enriched among these differentially expressed genes (Fig. 3B, Table S2). In both sets, the largest group of genes have been previously categorized as stress response genes (30% of the genes are differentially expressed in both comparisons, 15% of the genes in PA14 alone). As expected, genes related to metabolism were the next largest group of genes identified in both comparisons (15%, cf. 7% in PA14 alone). In comparison, genes related to proteolysis (7.6%) and extracellular material (6.6%) were the next largest group in those identified in PA14 alone (but only 2% and 1.7% among genes identified in both, respectively). This suggests that proteolysis and extracellular transport are processes involved in response to the virulence of PA14, rather than to the metabolic differences between *E. coli* and *P. aeruginosa*.

### Differential expression analysis using z11 as a control recapitulates known expression patterns, unveils expression changes in known infection-related genes, and discovers novel ones

Next, we sought to confirm that z11 is a more adequate/appropriate control for a PA14 infection by looking at the group of genes differentially expressed upon a comparison of PA14 with an OP50 control and a comparison of PA14 with a z11 control. The number of genes identified in both cases was similar, with a substantial overlap (Figure 3C). The fold-change in the expression of DEGs in each comparison was highly correlated (R^2=0.92, Figure 3D). This included a well-studied immune genes, *irg-1* (as well as *irg-2-3,-5,-6,-7*) ^23^; The TGF-beta transcription coactivator *sma-9* ^39^; as well as many lysozymes and C-type lectins ^40^.

We finally asked if the use of z11 as a control for PA14 studies increases the discovery power. To address this question, we focused on genes identified as differentially expressed on PA14 when using z11 as a control but not when using OP50. Among these 149 genes, 49 were categorized by WormCat as stress response genes, related to either detoxification (9 genes), pathogen response (9 genes), or others. These genes included *kgb-1, mlk-1,* and *vhp-01,* which encode a JNK-like MAPK, MAPKK, and MAPK phosphatase, respectively. While it is known that this pathway regulates the *pmk-1* MAPK pathway and therefore resistance to bacterial infection ^12^, the transcriptional induction of these genes was not previously observed in studies that used OP50 as a control.

The role of transcriptional regulators is also emphasized when using z11 as the non-pathogenic control. 39 genes that encode transcription regulators are identified in this analysis (among them eight nuclear hormone receptors, 17 other transcription factors, and 14 chromatin modification factors). Of these, 27 are up-regulated and 12 are down-regulated. In comparison, only 16 of these are found in an analysis that uses OP50 as a control.

Some of the transcription factors that are found with z11 control but not with OP50 control are of particular interest. The involvement of some of these genes in worm response to PA14 was previously discovered by other means. These include the bZIP transcription factor *zip-1*, the known regulator of *irg-1* ^24^, and the bZIP transcription factor *cebp-1,* which had been identified through mutagenesis analysis to affect worm survival of PA14 ^41^; The TFEB ortholog *hlh-30,* which regulates autophagy and aging ^42^, was known as a central regulator of worm immunity towards some pathogens including *E. faecalis* ^43^ and *S. aureus* ^44^, but its involvement in response to PA14 was not known; The CREB ortholog *crh-2* was known to regulate the expression of *mir-243* under PA14 pathogenesis, leading to diapause ^45^.

We also considered two transcription factors that were discovered in the z11 analysis and were not previously linked to PA14 infection: The gene *hlh-11* encodes a Helix Loop Helix transcription factor that has been recently shown to function in the intestine to modulate lipid metabolism in response to nutrient availability by activating the transcription of lipid catabolism genes to utilize fat ^46^; and the only worm FOXP ortholog *fkh-7,* which is expressed in the head, tail and intestinal neurons and is involved in the regulation of the transition to diapause (Chai, Cynthia, 2021). The expression of these two genes is significantly enhanced in the comparison between PA14 and z11 exposure, but not with OP50 (Fig. 4A). To test the significance of these transcription factors, we performed slow-killing assays on mutants of these genes and compared them with wild type N2. As above, all strains were grown on z11 for several generations before exposure to PA14. Both mutants showed a significantly reduced survival on PA14 lawn (Fig. 4B), suggesting that they play an essential role in worm response to PA14 pathogenesis.

**Figure 4.**
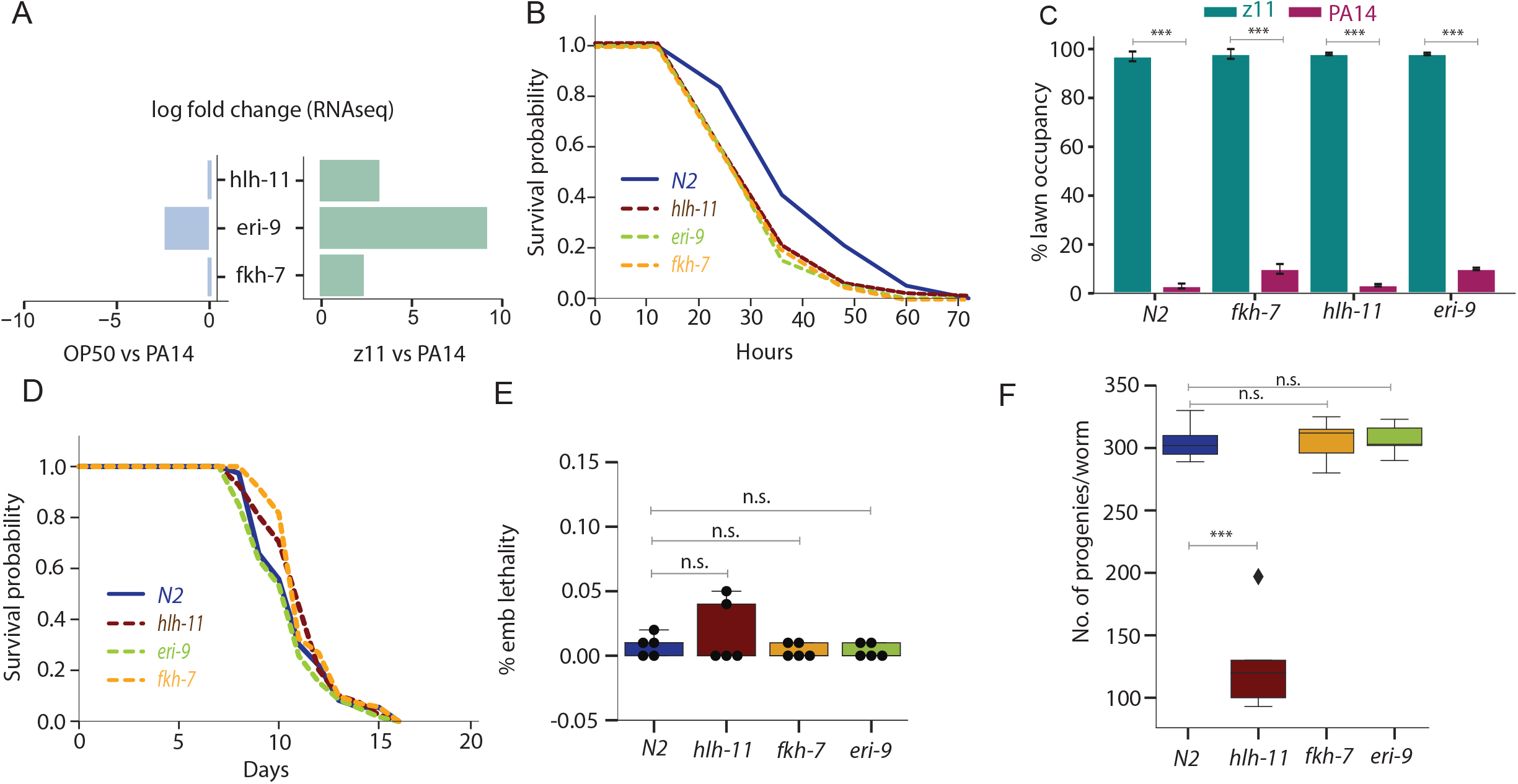
Using z11 as a control for differential expression analysis unveils new gene associated with host survival of PA14. (**A**) The expression of *hlh-11, fkh-7* and *eri-9* are significantly different between PA14 and z11 but not between PA14 and OP50. (**B**) Lifespan of *hlh-11(ok2944), fkh-7(ok455)*, and *eri-9(gg106)* mutants and wild-type N2 on z11 lawn at 25 degrees. P-values (relative to the wildtype): *hlh-11*(<0.54), *fkh-*7(<0.42), and eri-9(<0.397). (**C**) Lawn avoidance. P-values, *hlh-11*(<0.001), *fkh-7*(<0.001), and *eri-9*(<0.001). (**D**) Survival during PA14 slow killing assay. P-values: *hlh-11*(<0.009), *fkh-7*(<0.03), and *eri-9*(<0.008). (**E**) Embryonic lethality. P-values: *hlh-11*(<0.157), *fkh-7*(<0.198), and *eri-9*(<0.199). (**F**) Brood size for animals grown on z11. P-values, *hlh-11*(<0.001), *fkh-7*(<0.485), and *eri-9*(<0.432). The graphs are the combined results of 3 independent experiments. Each experiment included n = 30 adult animals per strain. *** p<0.001, n.s., non-significant.

Our list of differentially expressed genes included only three small non-coding RNAs. One of these genes, *eri-9,* was robustly up-regulated during PA14 exposure compared with z11 but was also induced in OP50. A killing assay of *eri-9* mutant worms also showed a decreased survival of PA14 (Fig. 4B), suggesting that this small RNA may play part in the regulation of response to PA14.

Finally, we asked if any of the reduced survival in these three mutants is due to inhibition of avoidance behavior, rather than molecular response. To address this question, we quantified their pathogen avoidance as described earlier and found that all strains were initially attracted to the PA14 lawn but altogether avoided the pathogen by 24 hours (Fig. 4C).

A reduced survival of mutant animals on a pathogen can be due to increased susceptibility, but it can also be due to unrelated processes that make the mutant less fit. For the three *hlh-11, fkh-7,* and *eri-9* mutants, however, we found no significant change in lifespan on the non-pathogenic z11 (Fig. 4D) and no increase in embryonic lethality (Fig. 4E), suggesting that these mutations do not make the animals generally unhealthy. In contrast, we find the number of progenies per animals is decreased in the *hlh-11* mutants (but not in the other two). Decreased fertility is typically associated with increased pathogen survival and does not in and of itself explains the observed increase in susceptibility to PA14.

## Discussion

Host response to a pathogen exposure is a multi-faceted process. Understanding this complex process and untangling its distinct aspects – metabolic adaptation, detoxification, innate and adaptive immune response, and behavior – benefits considerably from using a simple host. One advantage of working with a simple host model is the ability to carefully control the reference non-pathogenic diet, in a way that would minimize the impact on host-pathogen interaction during infection and simplify the interpretation of the results.

Here we show the benefits of a careful choice of a reference non-pathogenic species, which is more similar to the virulent pathogen in the well-studied model of intestinal infection of *C. elegans* by the opportunistic pathogen *P. aeruginosa* PA14. We show that this strain, *P. aeruginosa* z11, has no adverse impact on the worm and does not interfere with the virulence of PA14. Previously, attempts were made to use a mildly virulent PA14 *gacA* (a two-component regulator) mutant, which is required for the expression of virulence genes, including pyocyanin as a non-pathogen control for infection studies of PA14 ^24,29,48^. Although *gacA* mutants exhibited reduced pathogenicity, diminished *irg-1::gfp* induction, and decreased aversion behavior, compared to non-pathogenic *E. coli* strain, these outcomes are nonzero ^24,29,48^ due to failure to completely eliminate the PA14 pathogenicity. In contrast, z11 infection outcomes are comparable to the typical *C. elegans* laboratory diet of non-pathogenic OP50, suggesting z11 is better suited as a non-pathogenic control for PA14 infection studies.

We expected that with z11 as a control, fewer genes would be identified as differentially expressed upon infection by PA14. This effect turned out not to be very significant, suggesting that most of the C. elegans response to PA14 is – directly or indirectly – due to its virulence or the pathogenesis it causes.

In contrast, the use of z11 as a control allowed the identification of genes whose expression is induced but were undetectable when using the more distant *E. coli* strain as a control. Among these were genes whose link to PA14 response has already been identified by other means (genetic screens, association with other known factors, response to other pathogens) ^10,12,32^. Other genes previously unknown to be involved in response to PA14, and we confirmed that three of them – two encoding transcription factors and one small RNA – indeed impact worm survival of the pathogen. It would be of interest to learn more about the mechanisms behind these responses.

Many other pathogens – bacteria and fungi – have been used to study host response in *C. elegans*. Despite the practical advantage of using a single control in all studies, we propose that developing a specified within-species control for each pathogen could increase the discovery power.

## Materials and Methods

### Strains

*Caenorhabditis elegans* strains included in this work are N2 (wild type), VC241 *fkh-7(ok455)* X, RB2177 *hlh-11(ok2944)* III, and YY216 *eri-9(gg106)* III. Bacterial strains include *E. coli* OP50, *P. aeruginosa* z11, and *P. aeruginosa* PA14.

### Worm handing and Maintenance

Regular worm maintenance was performed as described earlier ^49^. Animals were age-synchronized by two methods: (1) hypochlorite treatment of gravid adult worms followed by collection and washing of the released eggs. Eggs were then allowed to hatch overnight in M9 buffer. (2) using a dissecting microscope to identify fourth larval stage (L4) hermaphrodites based on the size of the animal and the appearance of the vulva as a dark half circle.

### Lifespan assay

Age-synchronized young adult worms (30-40 worms per plate) were transferred to NGM plate seeded with appropriate bacterial lawn containing FUDR to prevent egg hatching. All lifespan analyses were conducted at 20 °C unless otherwise specified. Worm viability was determined every 1-3 days. A worm was scored as dead when it failed to respond to a gentle touch with a sterilized platinum wire. Any worms that were missing or having crawled off the plate were censored during analysis.

### Sterility and Fecundity assay

Sterility and fecundity assay was performed by individually plating synchronized L4 animal on NGM plates seeded with OP50 or z11. Embryos and hatched progenies were counted on day 1 to day 5 of egg laying. Eggs that failed to hatch after 2 days post egg laying were counted as dead. 10 fertile N2 young adult animals were individually cultured for 24 h, and their progeny were scored for sterility after growth for 3 days at 20 °C. Hermaphrodites were transferred to a fresh Petri dish after 2 days to eliminate self-progeny and transferred as necessary thereafter.

### Developmental assay

The synchronized L1 worms were transferred to NGM plates seeded with OP50 or z11 and assessed for L4 and young adult phenotypes. A dissecting microscope was used to identify fourth larval stage (L4) hermaphrodites based on the size of the animal and the appearance of the vulva as a dark half circle. Adult hermaphrodites were identified based on the fully developed vulva. The developmental stage of the worms was observed and quantified up to 46 hours post-transfer.

### Physiological assay

The physiological assay was started by placing L1 hermaphrodites on NGM plate seeded with OP50 or z11 and one young adult animals were examined using a dissecting microscope for the following phenotypes:

i. Thrashing: Worms from the test plates were washed and transferred to glass slide in a drop of M9 buffer. A 20-30 second movie of the motile worms was recorded and body movement in thrashing assay was assessed by observation. Thrashing movement was calculated by counting the rate of thrashing in M9 for 20-30 seconds as previously described ^50^. A thrash was defined as a head to tail sinusoidal movement.
ii. Pharyngeal Pumping: Worms were transferred from their population plate to the assay plates and left for 10 min to recover. Pumping was quantified by counting the number of grinder movements observed under stereo microscope contractions during a 20-sec interval ^51^. The pump rate was reported as pumps per minute.
iii. Body length and volume: To measure body length, images of the day 1 adult worms were taken. These images of the nematodes were skeletonized using ImageJ software. The length of the skeleton was used to determine the body length of the nematodes.

### Avoidance assay

Small lawns of z11, PA14, and OP50 were cultured on 6-cm NGM plates overnight at 37°C. Approximately 20 young adult animals grown on OP50 and z11 were put in the center of each bacteria lawn. The number of animals on each lawn was counted after 1-24 h ^26^.

### Slow killing assay

*P. aeruginosa* killing assay was done as described by Kirienko et al. ^52^. Briefly, overnight culture of P. aeruginosa PA14 strain was spread on Slow Killing (SK) media plates and incubated for 24 h at 37 °C and moved to 25 °C incubator. A total of 30–40 late L4 to young adult worms were transferred to seeded SK plate and a total of 100–120 worms (4 plates) were used per condition. An animal was classified as dead if it displayed no spontaneous movement and no responsive movement after prodding with a platinum wire. Lifespan (LS) was defined as the time from day zero to the last day of survival. The animals that escaped were excluded from the analysis.

### GFP reporter expression analysis

Induction of the expression of fluorescent reporter genes was measured as described ^24^. Overnight cultures of OP50, z11, and PA14 were seeded onto SK plates, incubated at 37 °C for 24 h, and then stored at 25 °C. Transgenic *irg-1::GFP* animals were transferred to respective bacterial lawn and incubated at 25 °C for 24 hours before washing and imaging a on glass slide using a Zeiss Inverted microscope at 10x magnification. Fluorescence intensity quantification was performed using ImageJ software.

### Statistical Analysis

Animal survival was plotted using Python matplotlib. The survival curves were considered significantly different from the control when the *p* values were <0.05. R (version 4.1.2) (Survival package) used to apply Kaplan-Meier method to calculate survival fractions and the log-rank test to compare survival curves. A two-sample Student’s *t* test for independent samples was used to analyze the other test, and *p* < 0.01 was considered significant. All experiments were repeated at least 3 times unless otherwise indicated.

### RNA-Seq and data analysis

Sample preparation: Infection assays were performed as described earlier. The bacterial strains were inoculated into 3 ml LB medium in a 15 ml polystyrene culture tube and grown overnight at 37°C. 200 μL of the overnight culture was spread on Slow-Killing agar medium in 10 cm Petri plates and incubated at 37°C for 24 h. We covered the entire SK plate with the bacterial lawn to decouple avoidance behavior from the immune response. After 24 hours at room temperature, each plate was seeded with approximately 2,000 synchronized young adult hermaphrodites. Plates were incubated at 25°C. The worms were collected at 12 hours post pathogen exposure and washed three times with ice-cold M9T buffer solution and three times with ice-cold M9 buffer solution by centrifuging at 2000 rpm for 45 seconds and then transferred to a 1.5ml tube for RNA extraction. Worm pellet was resuspended in TRI® Reagent (Sigma-Aldrich, St. Louis, MO) and stored at −80°C for TRI® Reagent (Sigma-Aldrich, St. Louis, MO) RNA extraction. According to the manufacturer’s instructions, total RNA was isolated in RNase-free water using Zymoclean RNA Clean & Concentrator™ (Zymo Research Corp.). Filtered high-quality reads were aligned to C. elegans genome WS281 (wormbase.org) using Kallisto with default parameters (version 0.46.2). For each sample at least 12 million reads were mapped to the C. elegans genome. Differential expression analysis was performed using the DESeq2 package (version 3.13) ^53,54^. GO enrichment analysis was performed by WormCat (http://wormcat.com/), a *C. elegans*-specific gene ontology enrichment analysis and visualization tool ^38^. For DEGs functional analysis, significantly up- and downregulated genes were input to WormCat using default settings and terms were considered statistically enriched when the reported p-adj values <0.01 conducted by Fisher’s exact test with Bonferroni correction.

## Acknowledgements

We would like to thank Alejandro Vasquez-Rifo and Victor Ambros for generous gift of *P. aeruginosa* z11 strain. This research was supported in part through NSF grant MCB-1946944. Some worm strains were obtained from the Caenorhabditis Genetics Center, which is funded by the NIH Office of Research Infrastructure Programs (P40 OD010440).

## Author Contribution

AR and EL designed the research. AR and EM performed the experiments. AR and EL provided mentorship, analyzed the data, and wrote the manuscript.

## Data Availability

Raw data presented in this manuscript have been deposited in NCBI’s Gene Expression Omnibus and are accessible through GEO (accession number TBD).

---

**Figure S1.**
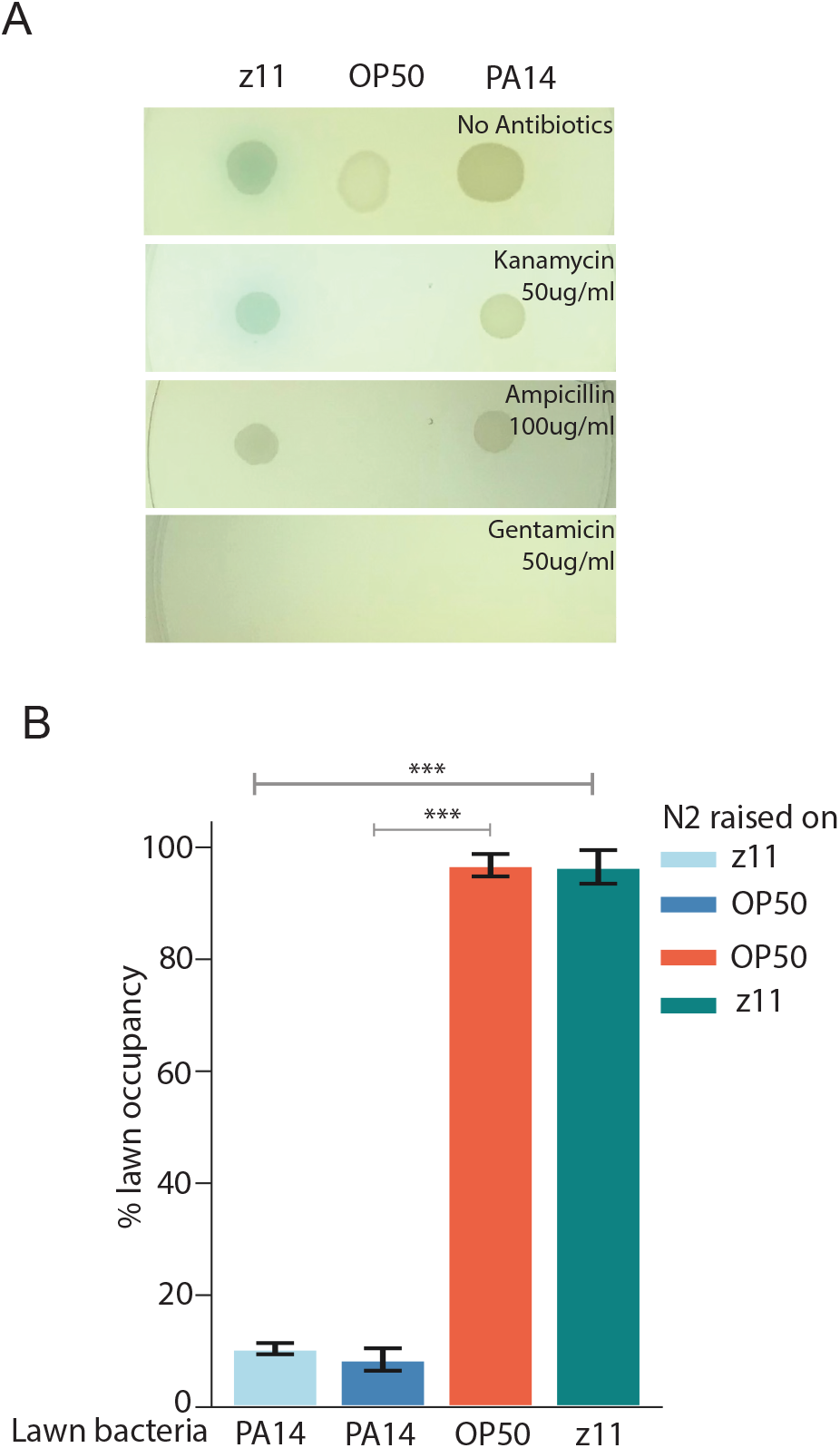
Properties of the lawns of the different bacteria. (**A**) Comparison of typical colonies of z11, OP50, and PA14 strains. All lawns were morphologically similar. Pyocyanin production by the *P. aeruginosa* strains (blue green pigmentation) can be seen around the colony. The two strains are naturally resistant to Kanamycin and Ampicillin. (**B**) Fraction of wild-type animals on lawns of PA14, OP50, and z11 bacteria lawn 24 hours after transfer to the lawn. ***: P-value < 0.001.

**Table S1.** Differentially expressed genes List of all differentially expressed genes between treatment comparisons (sheet 1: OP50 vs PA14, sheet 2: OP50 vs z11, and sheet 3: z11 vs PA14. count per million reads per gene values are shown for each treatment.

**Table S2.** GO enrichment categories. List of all enriched GO-term categories between treatment comparisons (sheet 1: OP50 vs PA14, sheet 2: OP50 vs z11, and sheet 3: z11 vs PA14. For each gene associated enrichment for GO categories are listed. Only terms with FDR < 0.01 and the most descriptive term for each unique gene are shown.

